# Protein inference using PIA workflows and PSI standard file formats

**DOI:** 10.1101/424473

**Authors:** Julian Uszkoreit, Yasset Perez-Riverol, Britta Eggers, Katrin Marcus, Martin Eisenacher

## Abstract

Proteomics using LC-MS/MS has become one of the main methods to analyze the proteins in biological samples in high-throughput. But the existing mass spectrometry instruments are still limited with respect to resolution and measurable mass ranges, which is one of the main reasons why shotgun proteomics is the major approach. Here, proteins are digested, which leads to the identification and quantification of peptides instead. While often neglected, the important step of protein inference needs to be conducted to infer from the identified peptides to the actual proteins in the original sample.

In this work, we highlight some of the previously published and newly added features of the tool PIA – Protein Inference Algorithms, which helps the user with the protein inference of measured samples. We also highlight the importance of the usage of PSI standard file formats, as PIA is the only current software supporting all available standards used for spectrum identification and protein inference. Additionally, we briefly describe the benefits of working with workflow environments for proteomics analyses and show the new features of the PIA nodes for the KNIME Analytics Platform. Finally, we benchmark PIA against a recently published dataset for isoform detection.

PIA is open source and available for download on GitHub (https://github.com/mpc-bioinformatics/pia) or directly via the community extensions inside the KNIME analytics platform.

## INTRODUCTION

Due to limitations in the measurable mass to charge range and in the resolution of mass spectrometers, measuring intact proteins at deep proteome level (i.e. detecting almost the complete proteome) is currently not possible. Therefore, shotgun proteomics using liquid chromatography coupled to mass spectrometry (LC-MS) is the state-of-the-art method to analyze peptides in a biological sample. Here, digested proteins are measured and much effort is used to identify as many spectra as possible, using lots of different search engines, and downstream analyze the proteins, using biological networks and pathways. The actual step of inferring the correct proteins ^1^ is rather neglected, though. While there are several tools and algorithms to perform the protein inference, which all have different restraints and assumptions ^2^, often protein inference is something thought of as automatically performed by the peptide search engines. This is only true for few search engines though, like X!Tandem and Mascot, and these can only employ peptide identifications found by the respective algorithms. Most of the widely used programs, e.g. MS-GF+ and Comet, return only the spectra identifications without any inference. An often employed approach to circumvent a more complicated protein inference is to allow only unique peptide identifications to infer proteins, i.e. peptides which are associated to exactly one protein in the protein database. This strategy is especially common in quantitative proteomics analyses. Another issue which should be addressed by a protein inference strategy is the creation of protein groups, instead of reporting protein representatives only. While this is done by most modern approaches, some older implementations still not address this thoroughly or hide the complete groups.

The tool *PIA – Protein Inference Algorithms*^3^ was introduced to facilitate the protein inference in an easy and user friendly way. PIA not only returns a comprehensive list of protein groups, but also shows the user the actual relations between peptide spectrum matches (PSMs), peptides and protein groups. The original publication highlighted the command line interface (CLI) and a web interface, as well as relatively simple KNIME ^4^ nodes, which in fact just wrapped the CLI commands. While the CLI can still be used for scripting and similar environments (e.g., where the user interface is not convenient), the web interface is currently no longer maintained.

Furthermore, PIA was integrated into other Java desktop applications and workflow environments such as the PRIDE Inspector Toolsuite^5^. The integration with PRIDE Inspector Toolsuite has enabled the quality assessment of thousands of complete submissions in PRIDE and is enabling the proper curation and review of the public proteomics data ^6^.

In addition, we have extended the features and support of KNIME nodes. KNIME is one of the leading workflow environments for data analysis and mining (https://www.knime.com/). With the KNIME Analytics Platform it provides an open source workflow infrastructure, which allows the creation and exchange of workflows not only in the field of proteomics, but in any data-driven science.

We have recently benchmarked PIA against other protein inference tools ^7^, and prove that it can accurately report the “correct” list of proteins. However, since the original publication, major features have been added to support PSI standard file formats and more peptide search engines. We will briefly recapitulate what PIA is for and how it works before highlighting the new features especially with respect to the KNIME nodes. Finally, we show an application example on a recent dataset for isoform characterizations, which is one main issue in protein inference and is far from being solved, as recently highlighted by the ABRF ^8^.

## USAGE AND FEATURES

### File format support

The input for PIA is the results of a peptide or spectrum search engine. PIA natively supports the identifications of Mascot ^9^ (Matrix Science Ltd., UK), X!Tandem ^10^ and the TXT output of Tide ^11^. Also the import of MSF files generated by ProteomeDiscoverer 1.3 and 1.4 (Thermo Fisher Scientific Inc.) is supported, including for example search results from Mascot, SEQUEST ^12^ and MS Amanda ^13^. Finally, the OpenMS ^14,15^ intermediate format idXML can be imported.

Originally,PIA supported the HUPO-PSI ^16^ standard formats mzIdentML ^17,18^. Recently,the import of files in the PRIDE database file format (PRIDE XML) was added as well as full support for the mzTab ^19^ file format, converting PIA in the only API (application programming interface) that can perform statistical analysis of proteomics data for all current PSI standards for protein identifications. For this reason, PIA is a key component of the PRIDE database ^20^ and is used to accurately assess the quality of the submitted peptide and protein evidences.

The support of PSI standard formats facilitates the import from almost all search engines, for which respective exporters or converters exist, as well as modern search engines which use the standard formats natively like MS-GF+ ^21,22^. As one principle, PIA does not perform any changes of the imported PSMs and the relations to the protein accessions reported by the search engine. If any peptide-protein mapping must or should be performed before the PIA analysis, this can be performed by external tools (e.g. the PeptideIndexer of OpenMS).

### Basic principles of a PIA analysis

An analysis using PIA is separated into several steps, a general workflow depicting these is shown in **Figure 1**. In the first step, the compilation, PIA structures the PSMs, peptides and proteins of the spectrum identifications into a directed acyclic graph, as described in ^3^. This is useful for all later analyses, as it speeds up the search of connected components, like PSMs belonging to a protein accession.

**Figure 1.**
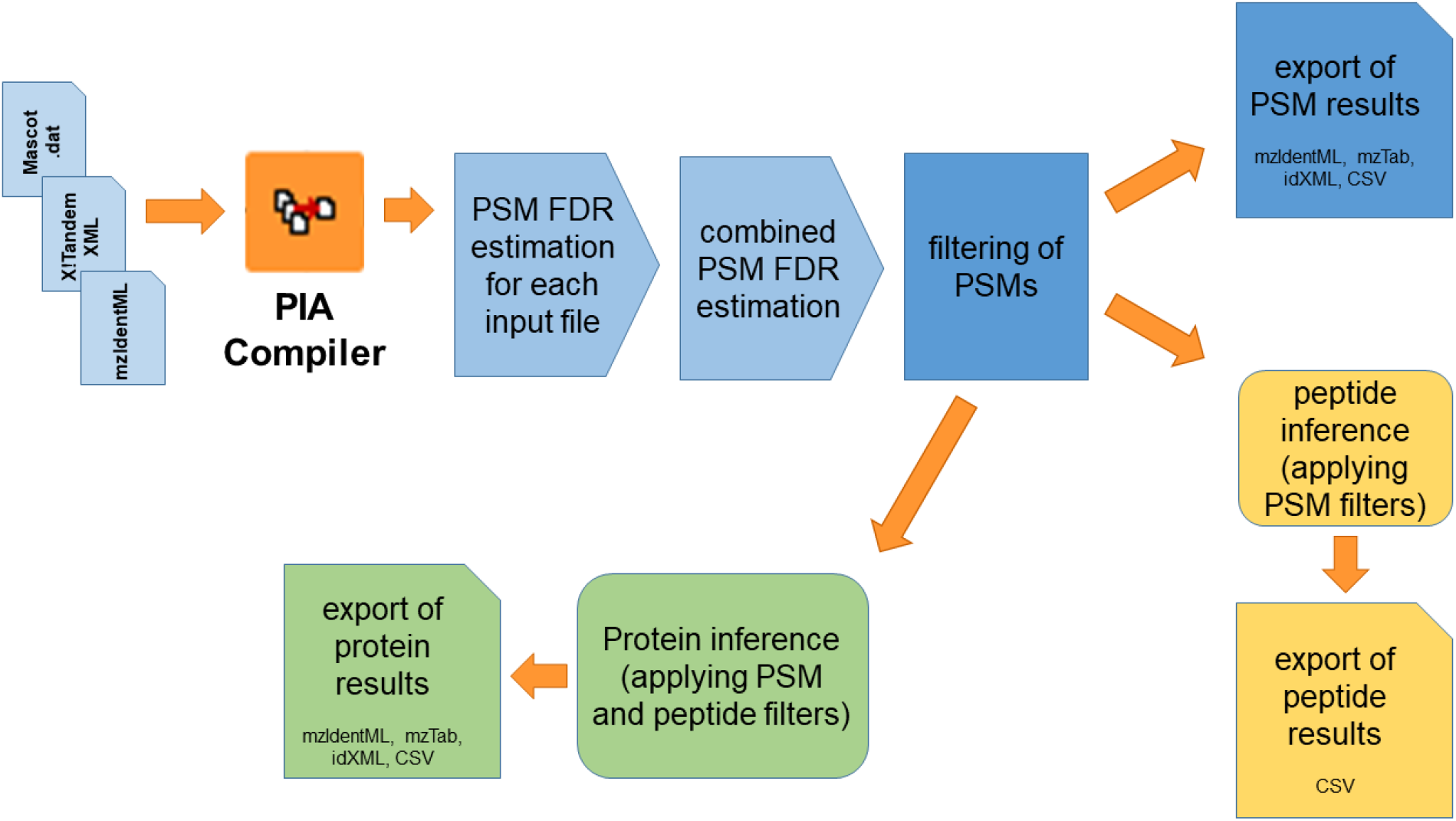
This chart is depicting a general analysis using PIA. First, the search engine results must be compiled into an intermediate format. Then the PSM FDR can be estimated for each file and the individual results can be combined. After filtering, a PSM level export can be performed. As the PSMs are the basis for the peptide and protein inference, both can be executed after all analyses on the PSM level are done. While the workflow only shows some of the supported input formats (for a more complete list refer to the text or the PIA tutorials), each level supports currently as output KNIME tables as well as the depicted (standard) file formats.

After the compilation is done, the actual protein inference analysis is performed. If multiple search results from the same MS run are to be analyzed, potential PSMs originating from the same spectrum can be combined into PSM sets. The false discovery rate (FDR) can be estimated using the standard target decoy approach ^23^ (TDA). Using this together with the *FDR Score* and *Combined FDR Score ^24^*, the peptide search results of multiple MS runs can be combined seamlessly. The PSM results, including the FDR estimation, of each file as well as of the combination of all files can be filtered and exported.

For the peptide level inference, PIA allows to filter the PSMs according to several properties like charge or measured mass error, as well as scores and FDR estimations. Identified modifications can be considered or left out for inference of peptides. In the latter case the peptide is defined be the amino acid sequence alone. Currently, it is not yet possible to select which modifications should be considered to distinguish peptides, like e.g. taking phosphorylations into account but ignoring oxidations of M.

PIA is based on the concept to always report protein groups, never just a representative. Hence, proteins or accessions, which have identical evidence on PSM and peptide level, are reported as one protein group (which might contain only one protein, though). Protein groups, which have overlapping but less evidence on PSM and peptide level as another group are reported as subgroups. For the protein inference, the user can select between three different methods, the *report all*, *Occam’s razor* and *spectrum extractor*. The first method just reports all possible protein groups, disregarding any sub-grouping. This option is only useful to scan an analysis for any specific identification, which otherwise might become a sub-group. The second option, *Occam’s razor*, is one of the most widely used parsimonious algorithms for protein inference. It reports a set of protein groups to explain all peptide evidences, which is as small as possible. *Spectrum extractor*, the default algorithm, is similar to *Occam’s razor*, but considers that some search engines report multiple PSMs per MS/MS spectrum. If this is the case, only one peptide per spectrum is assigned. Which peptide to assign is determined by the evidence of the protein without the respective spectrum. The search engine score or *FDR Score* of the PSMs which should be used by the inference algorithm to calculate the protein group score can be selected by the user, as well as any filters on properties (like mass deviation and charge) or scores. Also on the protein level the FDR can be estimated using the default TDA approach, as controlling the FDR on each level is highly recommended^25^.

All of the principles explained in the prior paragraphs are explained in more detail in the original PIA publication.

### Improved integration into KNIME workflows

The main change in the KNIME nodes of the original implementation is the change from a CLI wrapper, which was implemented using the generic KNIME nodes ^26^ (GKN), towards a more flexible native KNIME implementation of the nodes. While the GKN implementation needed a single node for each setting of a parameter or filter for the KNIME analysis, the new implementation has only two nodes, the *PIA Compiler* and the *PIA Analysis*.

The *PIA Compiler* is used to aggregate spectrum identifications from search engine results, create the acyclic graph and save it into an intermediate XML format. The node does not allow many settings and gives only some basic information about the input files, modifications and the total number of PSMs and peptides after the compilation.

The *PIA Analysis* node performs all other functionalities of PIA and lets the user adjust the settings at all levels of the analysis: the PSM, peptide and protein level as well as some general settings. The applied filters for each level can be adjusted and it can be selected whether the overview (or merge) of all files or the analysis of a specific input file should be reported. The reported lists of PSMs, peptides and protein groups are saved into KNIME tables and thus are directly available for all kind of other nodes, which allow further analyses inside the workflow environment, e.g. basic KNIME nodes for statistics or clustering as well as scripts for R ^27^ or Python.

As PIA allows a comprehensive analysis on PSM and peptide level besides the protein inference, the report on peptide and protein level can be disabled, if either is not required for the intended analysis. One of the new features is the ability to estimate the peptide-level FDR. Even if the peptide level FDR is currently not used by any of the protein inference algorithms, an analysis can be important. With this strategy PIA is used for the internal reanalysis of the PRIDE ^6,20^ database.

Besides the newly supported import formats mzTab and PRIDE XML, PIA supports also new export formats. To allow an easy exchange of results between scientists, mzTab is now supported. The idXML export for OpenMS compatibility was adapted to support OpenMS 2.x and the new protein grouping for quantification. For the deposition of results in an archive like PRIDE, it is recommended to use the standard format mzIdentML for data concerning MS/MS identifications. PIA now allows exporting into this format, using the recommended framing for describing the protein inference on all levels supported by this mzIdentML ^18,28^.

### Protein inference visualization

The main focus of PIA besides providing a comprehensive protein inference is to actually show the evidence behind the reason for reporting a certain protein group. This was one of the main aspects of the web interface and is now also available from the *PIA Analysis* node inside KNIME and as a Java generic component already in use in some proteomics tools (e.g. PRIDE Inspector Toolsuite^5^). **Figure 2** shows the visualization given by the *PIA Analysis* node, after the protein inference was performed. The respective view shows a list for the inferred protein groups with the respective scores, protein sequence coverages and target-decoy states, as well as lists for the peptide and protein levels, with all important information on these levels. Furthermore, the relations between the PSMs, peptides and accessions is depicted as a graph. This graph can give a comprehensive explanation for the causes, why a certain accession was selected in a protein group and others might be reported as a sub-group or not at all, using a color code: black bordered nodes are reported, while nodes without borders are in subgroups. The items of the currently selected protein group are intensely colored, while members of other groups are in pale shades. Finally, nodes which were available in the search engine results but filtered out, have no filling at all.

**Figure 2.**
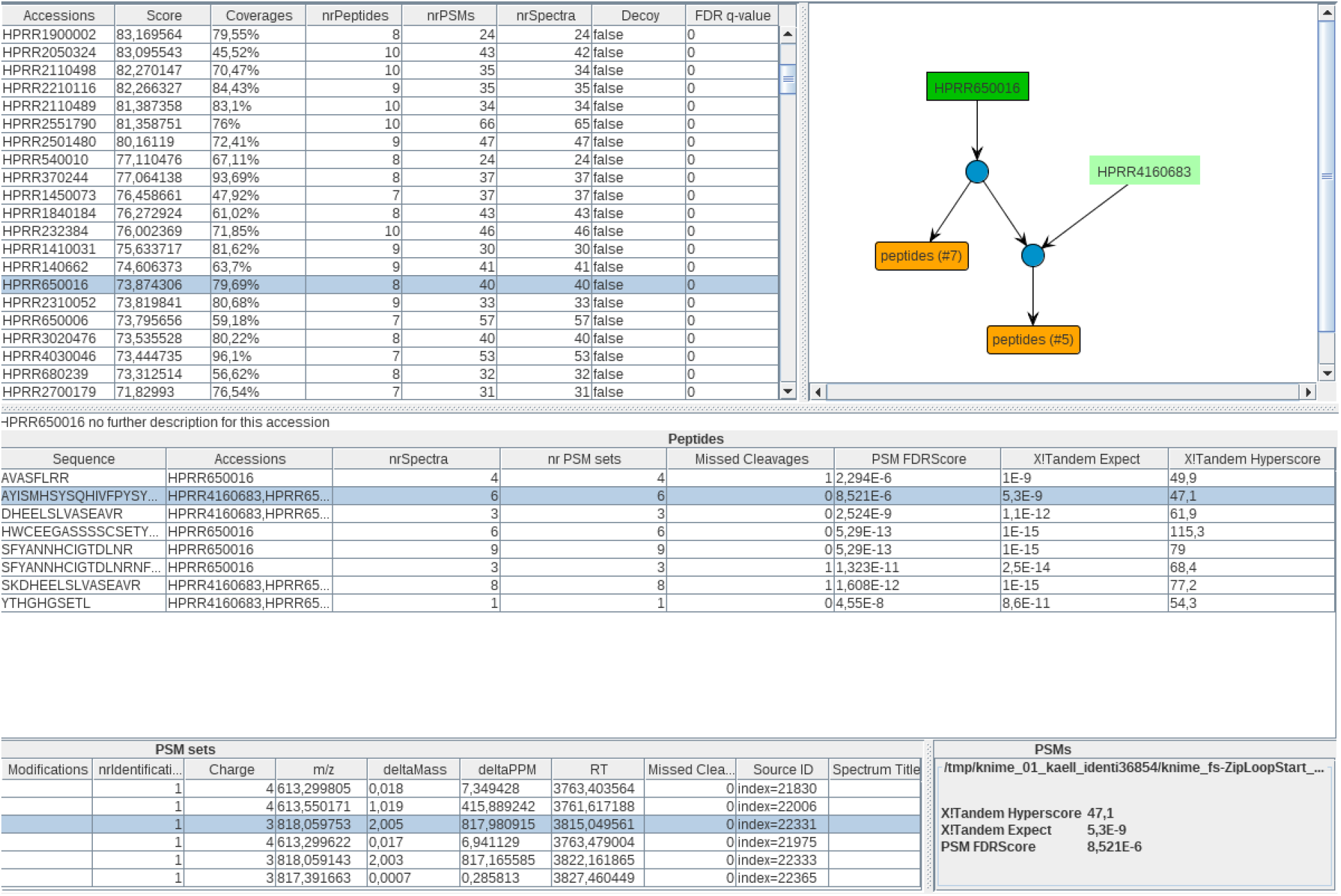
This screenshot shows the viewer of the PIA Analysis node. Lists of protein groups, the respective peptides and PSMs are given with all important information for each level. The relations between accessions, peptides and PSMs is reported in the graph on the upper right-hand side. The graph shows accessions (resp. proteins) in green boxes and peptides in orange boxes. The directed graph shows for each protein the contributing peptides. In the example, all peptides contribute to the upper group with the single accession HPRR650016, while only the lower peptides in the graph also contribute to HPRR4160683. Thus, that the protein group with the accession HPRR650016 is reported (green box with black border), while another protein group is reported with less evidence as a sub-group (green box without border).

Besides the viewer for the analysis results, also an optional spectrum viewer was implemented, using the ms-data-core-api^29^. This viewer (see **Figure 3**) does not visualize the actual annotation given by a search engine, but reconstructs the spectrum annotations given the actual spectra and the amino acid sequence together with any modifications. This viewer can only be used, if the MS/MS spectra are provided at the time of the PIA analysis.

**Figure 3.**
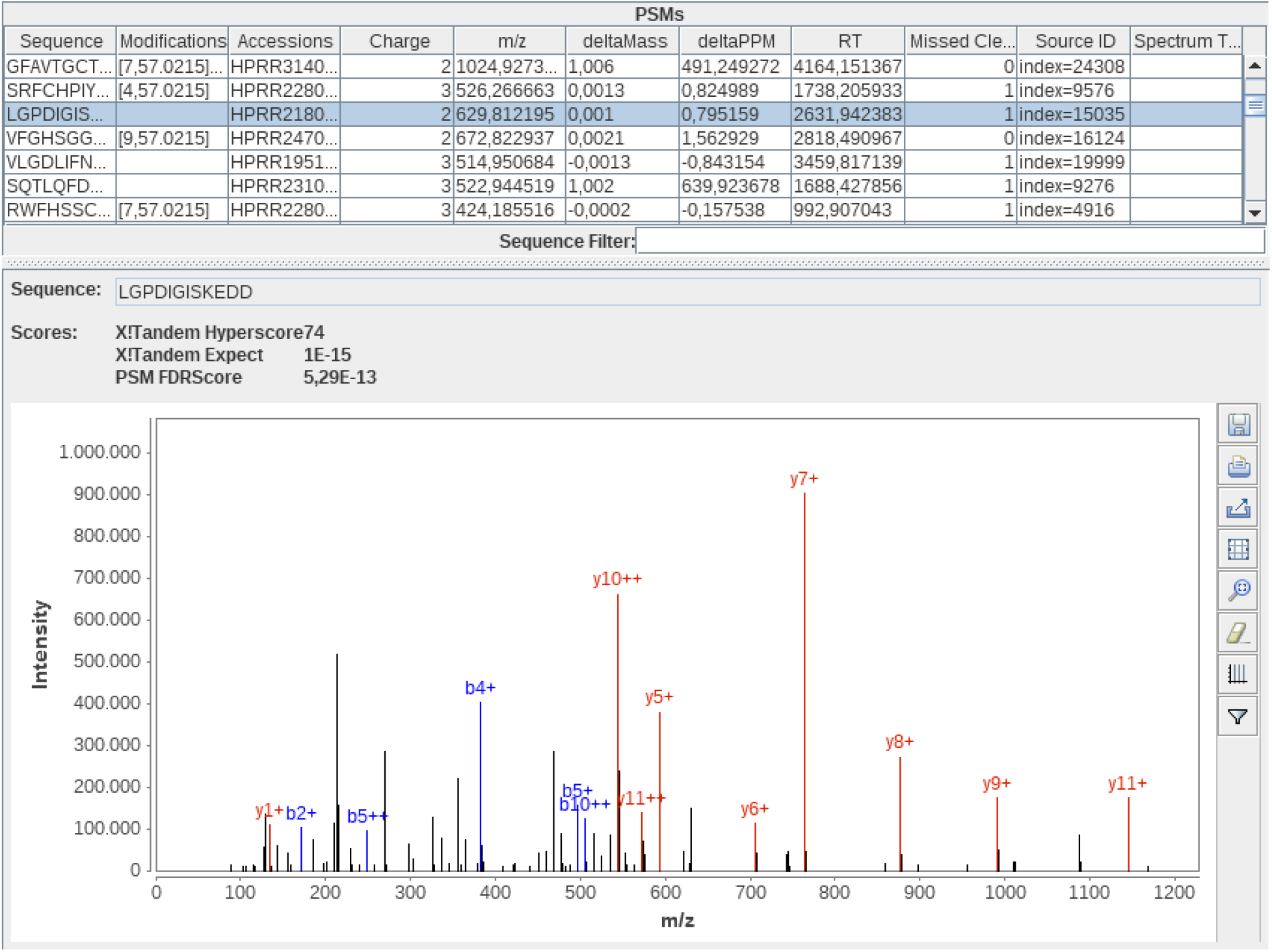
The spectrum viewer implemented in the *PIA Analysis* node.

### Application example: detection of protein isoforms

Recently, The *et al*. published a dataset^30^, which emulates homology and benchmarks the theoretical ability of different protein inference approaches to detect protein isoforms. In short, in this study several recombinant protein fragments (PrESTs) where expressed and analyzed using mass spectrometry, to simulate different levels of overlapping proteins. To test the performance of PIA, the data was downloaded from the PRIDE repository (accession number PXD008425). The raw files were converted using MSConvert^31^ and the vendor peak picking algorithm. The provided FASTA files were concatenated and a target decoy database was created by reversing the protein entries. All following steps, except the plotting, were performed inside a KNIME workflow (see supplemental data). The spectra were identified using X!Tandem with the following settings: 10 ppm precursor mass tolerance, 0.02 Da fragment mass tolerance, carbamidomethylation of C as fixed modification, no variable modifications and trypsin as cleavage enzyme. After the identification, the three replicates of each mixture given in the dataset (A, B and AB) were analyzed by PIA together, filtering the identifications on PSM level on 1% FDR. For the protein inference, the *Spectrum Extractor* was used with the *PSM FDR Score* as base score. The protein level results were finally used to execute the script provided by The *et al*., to plot the estimated q-values against the expected values. The resulting plots are given in the **Supplemental Figures S1-S3**. These figures show, that the PIA results are within the estimated bounds for a parsimony algorithm using product of PEPs, which is the category of PIA in the comparison, when compared to the plots given in the respective publication. **Table 1** shows the results of the analysis on the 5% protein-level entrapment and reported FDR. The table shows, that PIA slightly outperforms the test-implementation of the original manuscript, but only on the single mixtures, not on the combination of both. It must be considered, that the analysis was mainly focused on proteins and not protein groups though, and thus is biased against the main principles of PIA. Furthermore, for the analysis we performed with PIA a PSM FDR of 1% was used, as is recommended, and not a 5% threshold as was used in the prior publication.

**Table 1.** The number of inferred proteins at a 5% protein-level entrapment FDR and 5% protein-level reported FDR. This table shows the number of proteins, which were reported for the homology dataset by PIA and in parentheses the values expected by the original manuscript.

|  | 5% protein-level<br>entrapment FDR. | 5% protein-level<br>reported FDR |
| --- | --- | --- |
| mixture A | 187 (181) | 190 (191) |
| mixture B | 198 (196) | 209 (203) |
| mixture AB | 353 (360) | 358 (365) |

To show, how the PIA viewer can be used to further elucidate the reasoning behind some actual reported isoforms, see **Figure 2** and **Figure 4**. In Figure 2, the graph for the relations between accessions and peptides shows that a protein group containing the single protein HPRR2310052 is reported, which is correct for mixture A. HPRR4160683 on the other hand shares several peptides with the reported protein, but is not expressed itself in the given sample. Thus, there are no unique peptides found for the protein HPRR4160683 and the respective protein group was only reported as a sub-group. **Figure 4** shows a more complex example: here, the protein accessions HPRR2310052 and HPRR2310049 are reported in different protein groups, each one containing only one protein. For the accession HPRR3950112, which is actually not present in the given mixture, some spectra were wrongly identified in the sample, but filtered out by the FDR threshold (hence the corresponding peptides are filled in white in the figure), which finally resulted in the report of the two correct protein groups.

**Figure 4.**
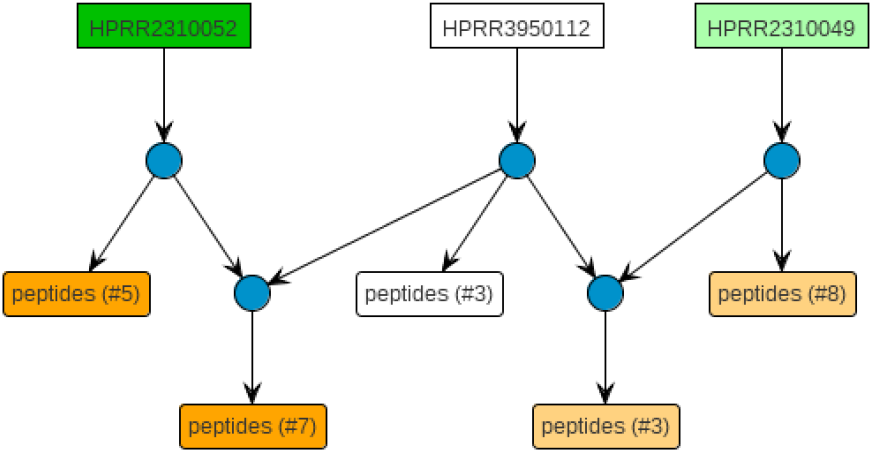
A more complex example of two correctly reported isoforms in the benchmark dataset. The proteins HPRR2310052 and HPRR2310049 are reported in different protein groups, containing only the one protein each, while HPRR3950112 is not reported and filtered out. The same color code as for Figure 2 applies. Furthermore, peptides which were identified in the original peptide search results but filtered out during the analysis, are depicted by a white background. Accessions, which lose all evidence due to this, are rendered in the same way.

## CONCLUSION

In this manuscript we highlighted some of the new features of PIA, especially the implementation inside the workflow and data analysis environment KNIME Analytics Platform, which is freely available and open source. This environment especially allows the repeated and automated execution of recurring tasks, but also the setup of complex analyses with subsequent annotations, statistics or machine learning after protein inference or quantification. These workflows can easily be exported and exchanged between scientists or can be deposited into archives like PRIDE to make the computational analysis as replicable as possible. PIA is currently the only protein inference tool for this environment, which also provides a graphical inspection of the results as well as KNIME tables as output. Furthermore, with the implemented support for native search engine results and HUPO-PSI standard file formats for spectrum identifications, a wide range of identification algorithms is available for a PIA analysis.

While PIA itself does neither use quantitative information for the analysis, nor reports any quantitative data, together with OpenMS it can help with the correct quantification of proteins, respectively proteins groups. OpenMS allows, besides many more features, the identification and quantification of peptides. It furthermore supports a tool for protein quantification, which can either perform a simple protein inference itself but also allows to import the inference information from other tools. For this, an idXML file containing protein group information is needed, which can be generated by PIA. OpenMS is also integrated into KNIME and thus PIA and the quantification nodes work together out of the box. Besides the mentioned methods to perform the protein inference using PIA, we also provide a Docker container via the BioContainers ^32^ group. This is especially useful for performing scripted or command line analyses in cloud environments.

PIA is open source under a three-clause BSD license and available for download via GitHub (https://github.com/mpc-bioinformatics/pia) or directly via the community extensions inside the KNIME analytics platform. Documentation and test data to start the usage of PIA is linked from the download location as well.

## Supporting information

## ASSOCIATED CONTENT

### Supporting Information

The following files are available free of charge at ACS website http://pubs.acs.org:

**Figure S1:** Plots created using the script of The *et al*. for mixture A.

**Figure S2:** Plots created using the script of The *et al*. for mixture B.

**Figure S3:** Plots created using the script of The *et al*. for mixture AB.

**Supplemental File 2:** KNIME workflow (knwf file) for the isoform analysis.

## Author Contributions

The manuscript was written through contributions of all authors. All authors have given approval to the final version of the manuscript.

## ACKNOWLEDGMENT

The development of PIA is funded by de.NBI (FKZ 031 A 534A), a project of the German Federal Ministry of Education and Research (Bundesministerium für Bildung und Forschung BMBF). ME is funded by PURE, a project of North Rhine-Westphalia, a federal German state.). Y.P.R is supported by BBSRC Grant ‘ProteoGenomics’ [BB/L024225/1], NIH ‘ProteomicsStandards’ Grant [R24 GM127667-01] and Two ELIXIR Implementation Studies. BE gratefully thanks Ruhr University Research School PLUS, funded by Germany’s Excellence Initiative [DFG GSC 98/3] for funding.

