## Supplementary material for "Protein inference using PIA workflows and PSI standard file formats"

**Table of content**:

- Overview
- Supplemental Figure S1: Plots created using the script of The *et al.* for mixture A.
- Supplemental Figure S2: Plots created using the script of The *et al.* for mixture B.
- Supplemental Figure S3: Plots created using the script of The *et al.* for mixture AB.
- Supplemental File 02: KNIME workflow (knwf file) for the isoform analysis

**Overview**:

This document contains plots using the data and script introduced in “A Protein Standard That Emulates Homology for the Characterization of Protein Inference Algorithms” by The et al (DOI: 10.1021/acs.jproteome.7b00899).

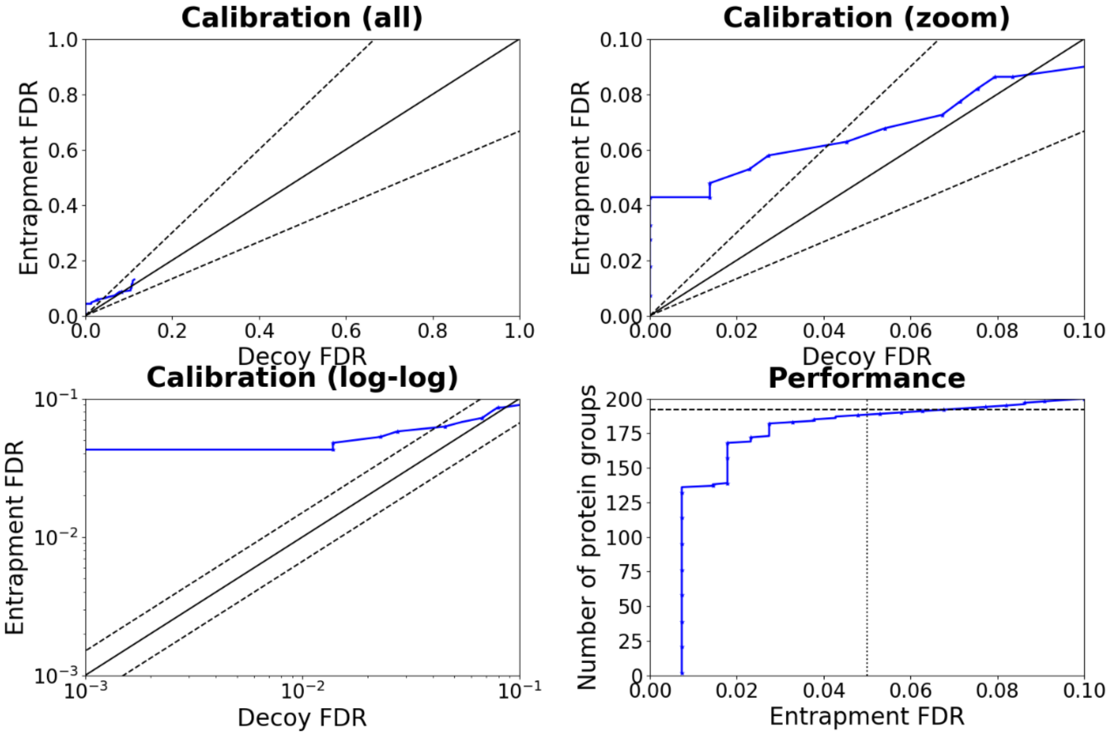


**Figure S1:** Plots created using the script of The *et al.* for mixture A.

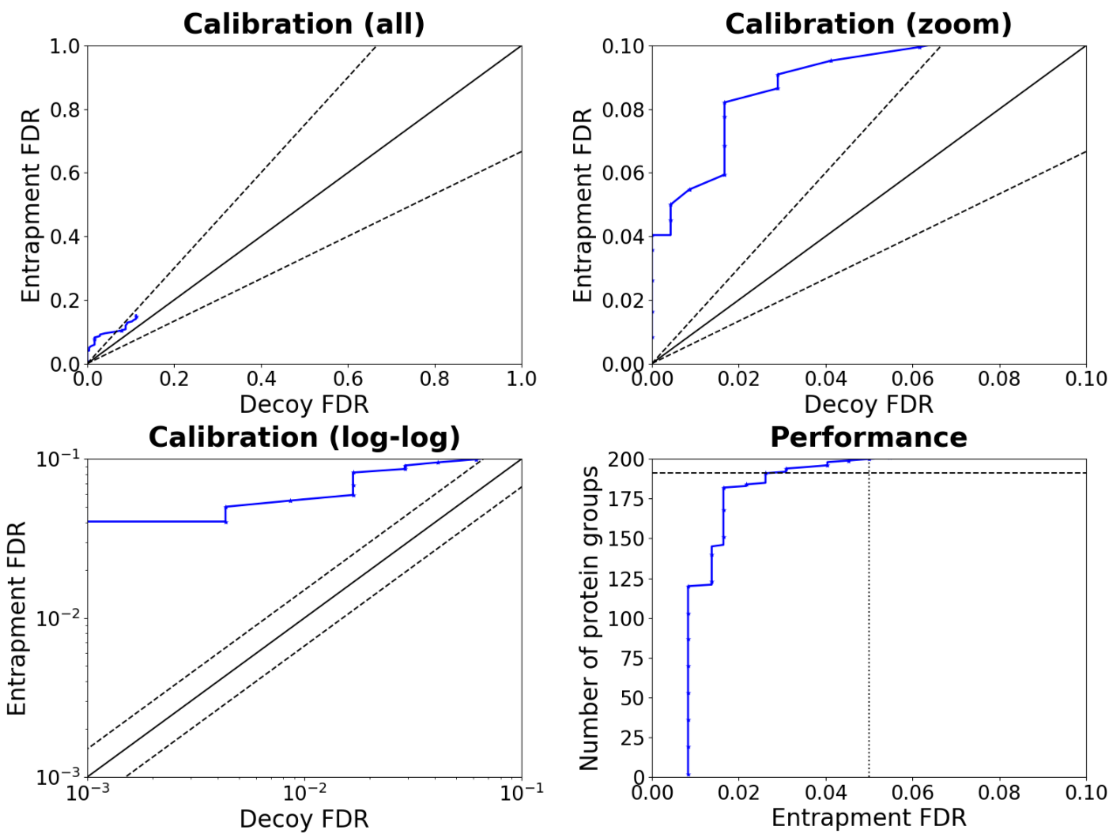


**Figure S2:** Plots created using the script of The *et al.* for mixture B.

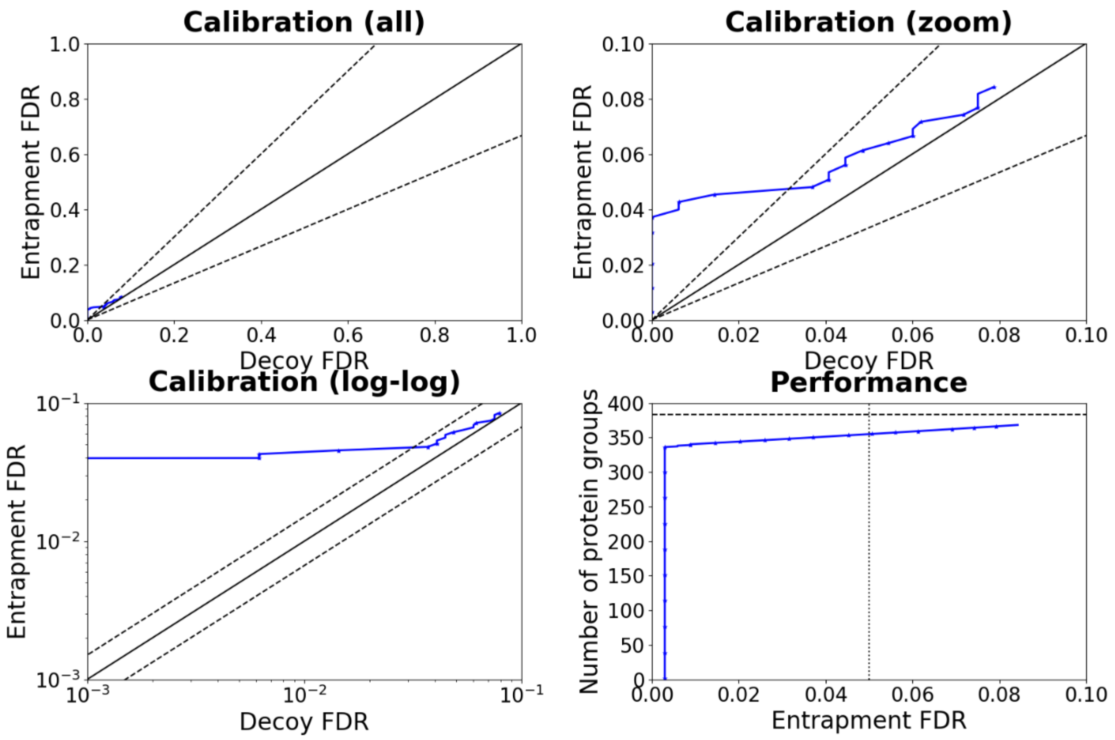


**Figure S3:** Plots created using the script of The *et al.* for mixture AB.
